# Sexually dimorphic traits and male-specific differentiation are actively regulated by Doublesex during specific developmental windows in *Nasonia vitripennis*

**DOI:** 10.1101/2020.04.19.048553

**Authors:** Yidong Wang, Anna Rensink, Ute Fricke, Megan C. Riddle, Carol Trent, Louis van de Zande, Eveline C. Verhulst

## Abstract

Sexually dimorphic traits in insects are rapidly evolving due to sexual selection which can ultimately lead to speciation. However, our knowledge of the underlying sex-specific molecular mechanisms is still scarce. Here we show that the highly conserved gene, *Doublesex (Dsx),* regulates rapidly diverging sexually dimorphic traits in the model parasitoid wasp *Nasonia vitripennis* (Hymenoptera: Pteromalidae). We present here the revised full *Dsx* gene structure with an alternative first exon, and two additional male *NvDsx* isoforms, which gives important insights into the evolution of the sex-specific oligomerization domains and C-termini. We show the sex-specific *NvDsx* expression throughout development, and demonstrate that transient *NvDsx* silencing in different male developmental stages dramatically shifts the morphology of two sexually dimorphic traits from male to female, with the effect being dependent on the timing of silencing. In addition, transient silencing of *NvDsx* in early male larvae affects male genitalia tissue growth but not morphology. This indicates that male *NvDsx* is actively required to suppress female-specific traits and to promote male-specific traits during specific developmental windows. These results also strongly suggest that in *N. vitripennis* most sex-specific tissues fully differentiate in the embryonic stage and only need the input of *NvDsx* for growth afterwards. This provides a first insight into the regulatory activity of *Dsx* in the Hymenoptera and will help to better understand the evolutionary and molecular mechanisms involved in sex-specific development in this parasitoid wasp, which can eventually lead to the development of new synthetic genetics-based tools for biological pest control by parasitoid wasps.

**Significance Statement:** In insects, male and female differentiation is regulated by the highly conserved transcription factor Doublesex (Dsx). The role of Dsx in regulating rapidly evolving sexually dimorphic traits has received less attention, especially in wasps and bees. Here, we mainly focused on Dsx regulation of two sexually dimorphic traits and male genitalia morphology in the parasitoid wasp, *Nasonia vitripennis.* We demonstrate that *Dsx* actively regulates male-specific tissue growth and morphology during specific developmental windows. These findings will help to better understand the molecular mechanisms underlying the rapid evolution of sexual differentiation and sexually dimorphic traits in insects, but may also be the starting point for the development of new tools for biological control of pest insects by parasitoid wasps.

## Introduction

Insects are well known for their distinct differences between male and female morphology. The reproductive organs and sexually dimorphic traits such as wing shapes and colour patterns are one of the fastest evolving traits in insects. Therefore, they are often used to distinguish species, as these traits can differ dramatically between species. Development of these fast changing sexually dimorphic traits usually depends on specific genes that are expressed in a tissue and sex specific manner (1). Many of these genes are under control of the transcription factor, Doublesex (Dsx), that initiates sexual differentiation in insects (2–4). *Dsx* pre-mRNA is alternatively spliced to yield male-specific *Dsx* (*DsxM*) or female-specific *Dsx* (*DsxF*) transcripts, resulting in distinct sex-specific protein isoforms (5). Two functional domains are present in all insect Dsx homologs: 1) the DM domain which consists of a DNA binding domain (DBD) and an oligomerization domain (OD1) with a zinc module and binding sites to facilitate binding downstream targets, 2) a dimerization domain (OD2) with a common N-terminal part and a sex specific C-terminal part. These C-terminal sex-specific differences are assumed to be involved in the regulation of the development of sex-specific traits (6,7). For instance, in *Drosophila,* DsxM represses and DsxF activates the expression of *bric-à-brac (bab)* which results in male-specific abdominal pigmentation (8). A similar action of Dsx is observed in the formation of sex combs, an important feature for performing mating rituals in males of some *Drosophila* species (9). In *Bactrocera dorsalis, Dsx* knockdown in females triggers the interruption of yolk protein gene expression, leading to ovipositor deformation and a delay in ovary development (10). It confirms the essential role of DsxF to initiate and maintain female development in Diptera. In Lepidoptera (butterflies and moths) and Coleoptera (beetles), sex-specific *Dsx* expression in both sexes also affects many sex specific traits. For example, transient expression of *DsxM* in female *Bombyx mori* silkmoths distorts the development of female-specific genital organs by partially introducing male characters in female genitalia (11), whereas introducing mutations in the male-specific *Dsx* isoform of *B. mori* yields either malfunctioning males or intersexes (12). In *Papilio polytes* butterflies, differential expression of *Dsx* isoforms results in sex-specific wing patterns (13). *Dsx* also promotes the mandible growth in males but constrains it in females of *Cyclommatus metallifer* beetles (14). In *Trypoxylus dichotomus* beetles, the formation of head horns is antagonistically regulated by *DsxM* and *DsxF*, while the thoracic horn formation is *DsxM* independent (15,16). Noteworthy, all these experiments were conducted in a specific developmental stage and only few focused on sexually dimorphic traits. However, the expression of *Dsx* starts in early embryogenesis and continues through maturity which implies its ongoing action (10,17,18).

Thus far, the specific role of *Dsx* in regulating sexually dimorphic traits has not been studied in the Hymenoptera. *Dsx* sex-specific splice variants and expression differences have been identified in honeybees, (*Apis mellifera*), bumble bees (*Bombus ignites),* Japanese ants (*Vollenhovia emeryi*) and fire ants (*Solenopsis invicta*) (19–23), but the role of *Dsx* in regulating reproductive organ development in males has only been shown in hymenopteran sawflies, *Athalia rosae* (24). To understand the rapid evolution of sexual dimorphism in insects and to compare the molecular mechanisms of *Dsx*-mediated regulation, it is paramount to gain more knowledge on how *Dsx* directs sex-specific and sexually dimorphic trait development in the Hymenoptera. Therefore, we set out to 1) identify the sex-specific traits that are regulated by *Dsx* and 2) study the timing by which *Dsx* regulates these traits during development in the parasitoid wasp *Nasonia vitripennis*.

*Nasonia vitripennis* belongs to the genus *Nasonia* (*Hymenoptera: Pteromalidae*), and has become the model system for parasitoids as it is highly suitable for evolutionary and developmental genetic studies. Multiple sexually dimorphic traits within the genus *Nasonia* make it a perfect system to study sex differentiation (25,26). For instance, the femur of the hind legs and the antennae of female *Nasonia* are pigmented with a dark-brown color, whereas in males these are pale yellow (26). Only in *N. vitripennis,* males have dramatically shorter forewings and are flightless (26), while the females can fly and have large forewings in all species. Although *Dsx* has been identified in *N. vitripennis* (27), its role in regulating these sexually dimorphic traits has not been shown before. Previous research identified one male *Dsx* (*NvDsxM*) and one female *Dsx* (*NvDsxF*) splice variant (27) and showed that the conserved splicing factor Transformer (Nvtra) is responsible for female-specific splicing of *Dsx* transcripts, while in males the *Dsx* transcript is spliced by default into the male isoform (28).

To assess the role of *Dsx* in regulating sexual dimorphism in *Nasonia,* we first revisited the *NvDsx* genetic structure. We confirmed existing splice-variants and identified an alternative starting exon and two additional male *NvDsx* splice-variants. We then used RNA interference to temporally silence *Dsx* in different male developmental stages to study its function in sexual differentiation and dimorphism. Ultimately, our results provide a great start to study the molecular basis of sexual differentiation and rapidly evolving sexual dimorphism in insects in general and particularly in the Hymenoptera.

## Results and discussion

### Identification and characterization of two additional male-specific *Dsx* splice variants

Previously, only one female (*NvDsxF*) and one male (*NvDsxM*) *Dsx* splice form were identified in *N. vitripennis* (27). The *NvDsxF* splice variant shares the same first four exons with the *NvDsxM* splice variant, but 108 bases are spliced out from exon 5 which leads to female-specific exons 5F and 6F. In *NvDsxM,* exon 5M results from retaining this 108bp intron, a form of alternative splicing which has not been observed for *Dsx* in other insects (27) (Fig. 1, *NvDsxF* and *NvDsxM1*). This splice form translates into a Dsx protein with a very short OD2 domain of only four amino acids (aa). The gene structure of this published *NvDsx* does not resemble any other published *Dsx* structures. To investigate the possibility of additional splice variants of *Dsx* pre-mRNA in *N. vitripennis,* we performed extensive 5’ and 3’ RACEs. The results reveal a novel alternative starting exon that we designated exon 1’ (Fig. 1). Note that the start codon of protein translation is in exon 2. Using RT-PCR to target exon 1’ to exon 5 we verified its presence in transcripts of both sexes but with a higher frequency in female transcripts (Fig. S1). Alternative splicing at the 5’ end has not been shown for *Dsx* before. We assume that the alternative exon is used in all splice variants (Fig. 1), although long-read RNA sequencing would be required to confirm this. From here on we refer to the alternative variants collectively. The 3’ RACEs identified two additional longer male splice variants (Fig. 1). We designated the previously published male-specific variant *NvDsxM1* (M1: NM_001162517; M1’: MT043363); the newly found longer male splice forms were designated *NvDsxM2* (M2: MT043359; M2’: MT043360) and *NvDsxM3* ((M3: MT043361; M3’: MT043362). All male isoforms share the same exons 1 (or 1’) to 5 with the female isoform *NvDsxF* (F: NM_001162518; F’: MT043364). The *NvDsxM2* open reading frame ends with a stop codon in exon 8 and translates into a protein of 328 aa which is 105 aa longer than the open reading frame of *NvDsxM1*. Compared to *NvDsxM2,* we find that exon 8 is spliced out in *NvDsxM3* which puts the stop codon in exon 9 and yields a protein of 330 aa. These two additional male specific splice variants skip the female-specific exon 6 completely which is a shared feature of male splice forms in many insect species including fruit flies (*D. melanogaster*) (29) (Fig. 1), wild silkmoths (*Antheraea assama*) (30), honeybees (*A. mellifera*) (21) and fire ants (*S. invicta*) (20). We did not find any non-sex specific splice variants in *N. vitripennis* as opposed to *A. mellifera* (Fig. 1) (21). NvDsxM2 and NvDsxM3 protein isoforms are almost completely identical and only differ at the C-terminus after the male-specific OD2 domain (Fig. S6A).

**Figure 1.**
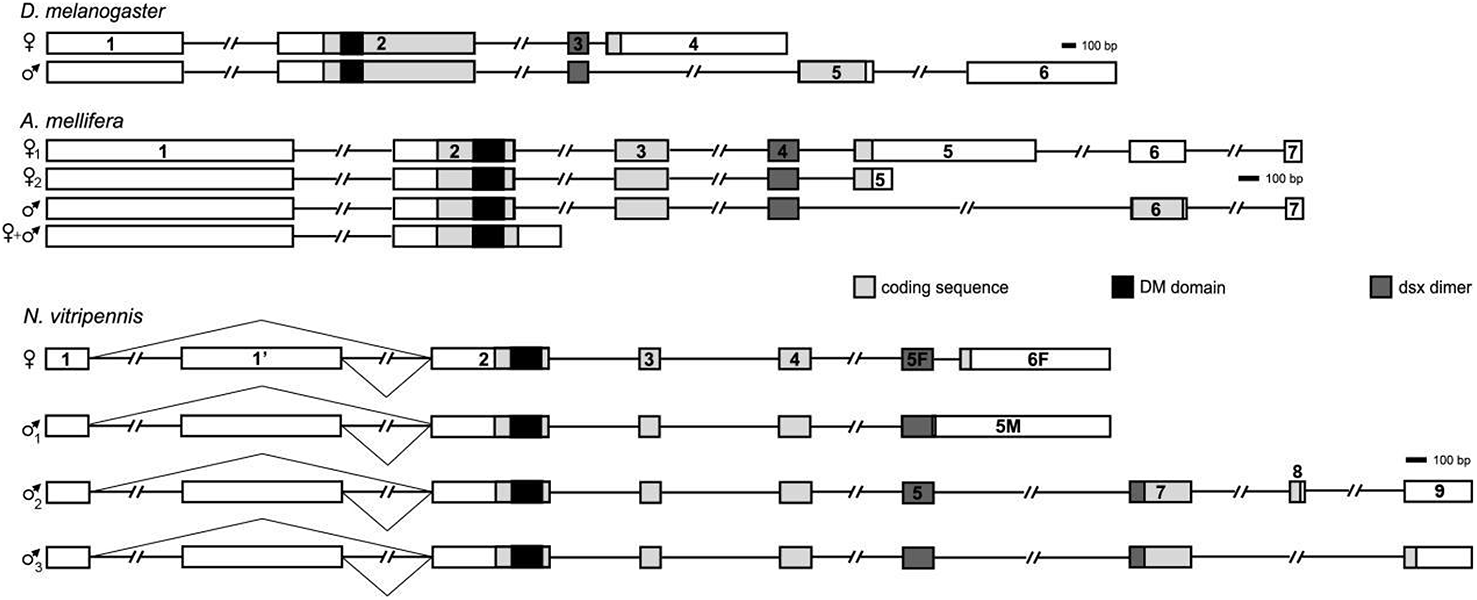
Comparison of insect *Doublesex* (*Dsx*) gene models. The eight *N. vitripennis* male and female *Dsx* splice variants are compared to *Dsx* splice variants of *D. melanogaster* and *A. mellifera*. Figure was modified from Oliveira *et al.* (27) to include the alternative 1^st^ exon (1’) and the additional *NvDsx* male splice variants. Male and female specific splice forms are indicated by ♂ and ♀. Exon numbers are noted on each exon. Light grey, black and dark grey blocks indicate coding region, DM domain and OD2 Dsx dimer region respectively. Scales are provided behind each species splice structures.

### *NvDsxM2* and *NvDsxM3* are spliced by default

To determine whether the additionally identified splice variants are exclusively expressed in males, we first observed the overall expression of different *Nvdsx* splice forms in both males and females. Across development, *NvDsxF* is only expressed in females while *NvDsxM1, NvDsxM2* and *NvDsxM3* are highly expressed in males (Fig. S2), and expressed at a low level in females. Only in males a slight increase of *NvDsxM1,* −*2 and* −*3* expression from 36h toward the early pupal stage was observed. Afterwards, the expression remains relatively constant until adult stage (Fig. S2). Overall, *NvDsxM2* is more abundant than *NvDsxM3* (Fig. S2). Females express *NvDsxM1*, *NvDsxM2* and *NvDsxM3* at a low level probably due to leaky default splicing of male-specific transcripts (Fig. 2B and Fig. S2).

**Figure 2.**
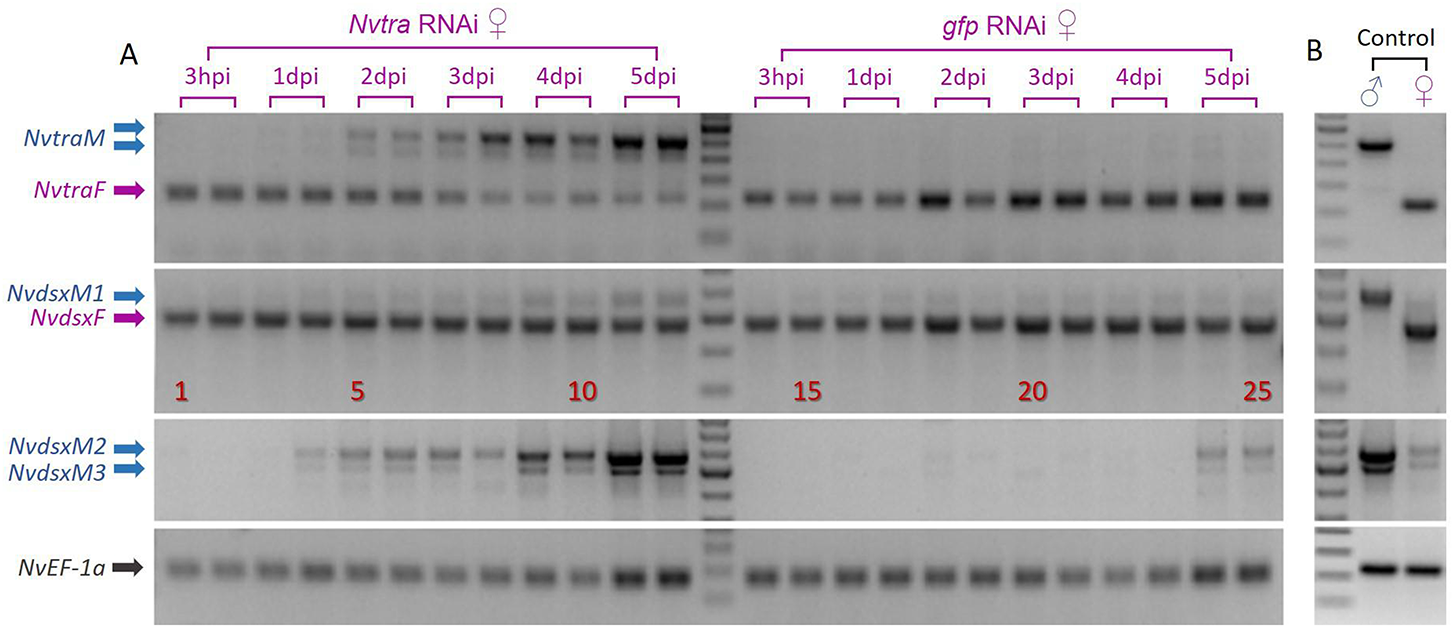
Confirmation of sex-specific *Dsx* splicing after *Nvtra* knockdown in *N.vitripennis.* (A) Sex-specific splice variants of *Nvtra* and *NvDsx* in *N. vitripennis* females that are injected in the 4th instar larval stage with *Nvtra* dsRNA or *gfp* dsRNA. (B) Male and female non-treated (NT) control samples for each primer sets. Arrows indicate male (in blue) and female (in purple) splice forms of the different genes. Lanes 1-12: Female *Nvtra* knockdown samples from 3-hour post injection (hpi) to 5-day post injection (dpi). Lane 13: 100bp ladder (Thermo). Lanes 14-25: Female *gfp* knockdown samples from 3dpi to 5dpi. For each stage two samples are used. *NvEF-1a* is used as expression and loading control.

We have shown previously that parental *Nvtra* RNA interference (RNAi) in the female pupal stage leads to a shift in splicing from *NvtraF* to *NvtraM* (28). This subsequently induces a shift from *NvDsxF* to *NvDsxM* (now *NvDsxM1*) splicing in both the treated adult females and the offspring, without consequences for adult female fertility (28). The reduced levels of NvtraF are no longer sufficient to regulate the splicing of all *Nvdsx* transcripts into the female-specific form, leading to default expression of *NvDsxM1* splice variants. To verify the default and NvTra independent splicing of *NvDsxM2* and *NvDsxM3*, we used *Nvtra* RNAi in the last larval instar (4^th^ instar) of females. RT-PCR was applied to amplify the different *NvDsx* and *Nvtra* splice variants from 3 hours post injection (hpi) to 5 days post injection (dpi). After confirming that *Nvtra* RNAi leads to *NvtraM* splicing in females (Fig. 2), we did not observe a clear reduction of female *NvDsxF* expression, but a minor increase of *NvDsxM1* expression from 4dpi onwards was observed. *NvDsxM2* and *NvDsxM3* expression levels increase substantially from 1dpi onward compared to *gfp* RNAi control (Fig. 2).

### NvDsxM represses female-specific pigmentation

To assess the sex-specific function of Dsx, we studied the phenotypic effect of *NvDsx* silencing at different developmental time points. We were unable to silence the male-specific splice variants of *NvDsx* specifically (Fig. S3B), therefore we targeted the common region of *NvDsx* (*NvDsxU*). We observed that the developmental stage at which *NvdsxU* dsRNA was injected greatly influenced the expression levels of *NvDsx* measured using qRT-PCR. We found no significant decrease of *NvDsx* expression in the 2^nd^ instar *NvDsx* knockdown (*NvDsxU*-i) male larvae in comparison to non-treated (NT) and *gfp* mock (*gfp*-i) when sampling after the white pupal stage (Fig. S3*A*). The strong increase in *NvDsx* expression from 4^th^ instar male larvae onwards (Fig. S2) may be the main cause as the limited amount of siRNA present in the 2^nd^ larval instar is not capable of silencing the sudden increase of transcript numbers in later developmental stages. However, this allows us to transiently suppress *NvDsx* during different developmental time windows. Injecting *NvDsxU* dsRNA in 4^th^ instar male larvae led to a significant reduction of *NvDsx* in newly emerged adults when compared with both NT (Tukey’s HSD, P<.001) and *gfp*-i males (Tukey’s HSD, P<.001) (Fig. S3*B*). It confirmed a successful *NvDsx* knockdown starting in 4^th^ instar male larvae with an effect lasting into adulthood. The 4^th^ instar *gfp*-i males also showed significant reduction of *NvDsx* expression compared to NT (Tukey’s HSD, P=0.021) suggesting that in the 4^th^ instar larvae there might be a non-target effect of *gfp* dsRNA as was shown before in the honey bee (31) (Fig. S3*B*). Clear reduction was observed at the adult stage of pupal stage *NvDsxU*-i males when compared to NT (Tukey’s HSD, P=0.003) and pupal stage *gfp*-i males (Tukey’s HSD, P=0.002) (Fig. S3*C*) which indicates that we also obtained a highly significant knockdown of *NvDsx* from the pupal stage onwards.

Non-treated and *gfp*-i males showed unpigmented antennae and legs (Fig. 3*A, B, C*), whereas females have pigmented antennae and legs (Fig. 3*D*). *NvDsx* knockdown (*NvDsx*-i) in 2^nd^ instar male larvae increased the amount of pigmentation in the adults slightly (Fig. 3*F*), but depleting *NvDsx* from 4^th^ instar male larvae onwards increased pigmentation in their adult stage to the point that it resembled female-like pigmentation (Fig. 3*E*). No pigmentation was observed in adult males when *NvDsx* was depleted from the early white pupal stage onwards (Fig. S4).

**Figure 3.**
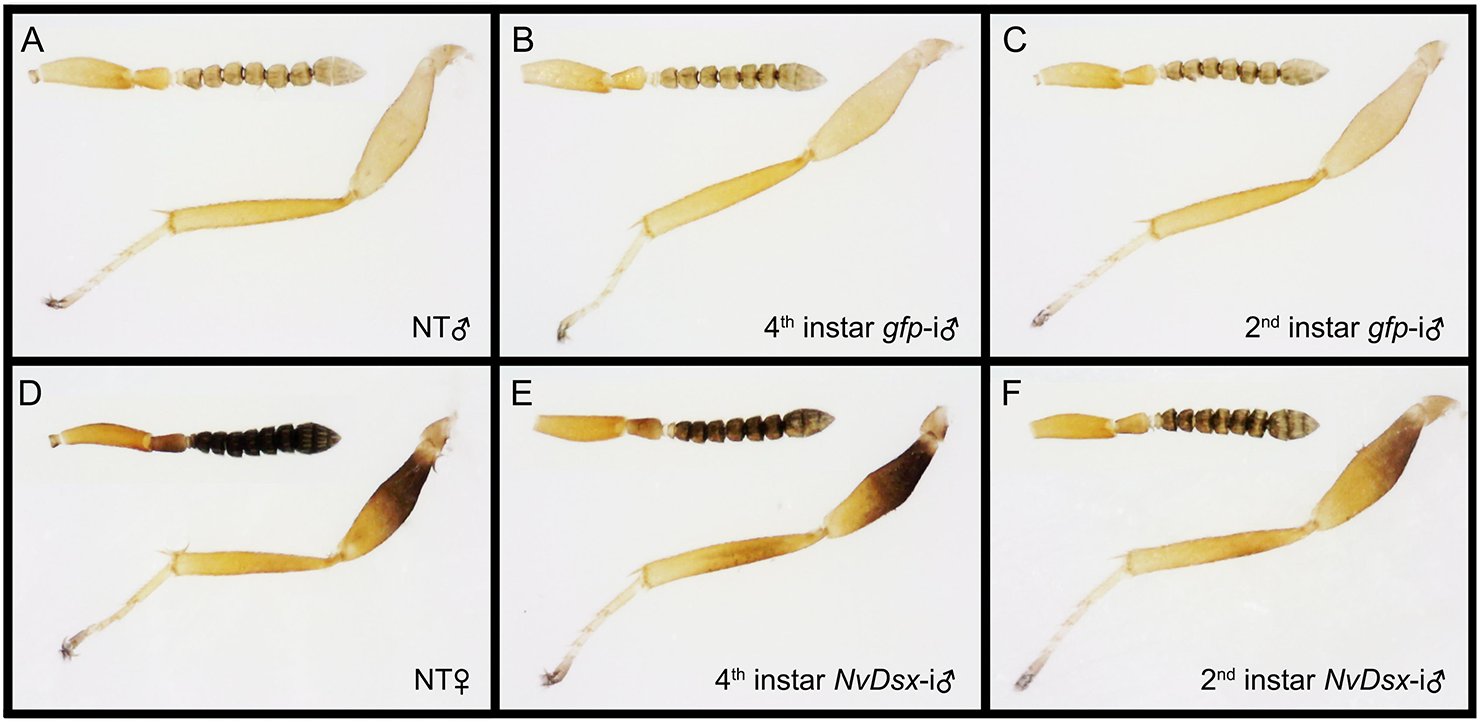
Pigmentation of antennae and hind legs of adult wasps after *NvDsx* or *gfp* dsRNA treatment in different developmental stages. (A) non-treated (NT) male (B) 4th instar *gfp* RNAi (*gfp*-i) male (C) 2^nd^ instar *gfp*-i male (D) NT female (E) 4^th^ instar *NvDsx* RNAi (*NvDsx*-i) male and (F) 2^nd^ instar *NvDsx*-i male. Light source and exposure time are equal for all pictures.

We did not find changes in pigmentation intensity by *NvDsx* RNAi experiments in 4^th^ instar female larvae (Fig. S5), and silencing *NvDsx* led to female-like pigmentation in males. Therefore, we conclude that female pigmentation is the default developmental state, and in males NvDsxM actively interferes with the melanin synthesis pathway. In *D. melanogaster,* expression of sex-specific abdominal pigmentation is regulated by interaction of the HOX protein Abdominal-B (Abd-B) and Dsx. Combined input of Abd-B and DsxF leads to the expression of *bric-a-brac* (*bab*) which suppresses the pigmentation in the posterior abdomen segments while DsxM represses *bab* to maintain the posterior abdomen pigmentation (8). As we found no true intersex phenotype, we hypothesize that NvDsxF plays no role in enhancing expression of the genes required for pigmentation in females. The slight reduction in antenna pigmentation of 4^th^ instar *NvDsx*-i males (Fig. 3E) is likely due to 1) a structural difference in the antennae with males having less plate organs on the sub-segments than females causing a different color reflection (32), and/or 2) incomplete knockdown of *NvDsx.* The deposition of the pigments in the cuticle often starts from late pupal to early adult stages in *Drosophila* (33). Therefore, initiating the RNAi response from 4^th^ instar larva onwards creates enough time to deplete *NvDsx* mRNA and NvDsx protein production to activate the expression of pigmentation genes in *N. vitripennis* males.

### *NvDsx* silencing leads to increased forewing size of males

The second sex-specific character that we assessed was forewing size and shape in *N.vitripennis* adult males. Silencing *NvDsx* in both 2^nd^ and 4^th^ instar larvae resulted in increased forewing length and width when compared to *gfp*-i (Tukey’s HSD, P<0.001) and NT males (Tukey’s HSD, P<0.001) (Fig. 4*B* and *C*). Length/width (L/W) ratio of *NvDsx*-i treatments also showed severe reduction compared to *gfp*-i (Tukey’s HSD, P<0.001) and NT males (Tukey’s HSD, P<0.001) (Fig. 4*A*) indicating that the forewing became wider and more female-like. However, the forewings of the *NvDsx*-i males were still smaller and narrower than those of the NT females (Length: Tukey’s HSD, P<0.001; Width: Tukey’s HSD, P<0.001; L/W ratio: Tukey’s HSD, P<0.001) (Fig. 4*A, B* and *C*). Interestingly, here we found an effect of timing as the forewing length and width increased significantly between 2^nd^ and 4^th^ instar larvae *NvDsx*-i treatments in which earlier stage *NvDsx* knockdown led to more severe changes (Tukey’s HSD, P<0.001), steering the L/W ratio towards NT females dramatically (Tukey’s HSD, P<0.001). No significant differences were observed between 2^nd^ and 4^th^ instar *gfp*-i males (Tukey’s HSD, P=0.062), 2^nd^ instar *gfp*-i and NT males (Tukey’s HSD, P=0.070) and 4^th^ instar *gfp*-i and NT males (Tukey’s HSD, P=0.999) (Fig. 4*A, B* and *C*). Depleting *NvDsx* from the white pupal stage onwards had no effect on forewing size or shape in males (Fig. S4).

**Figure 4.**
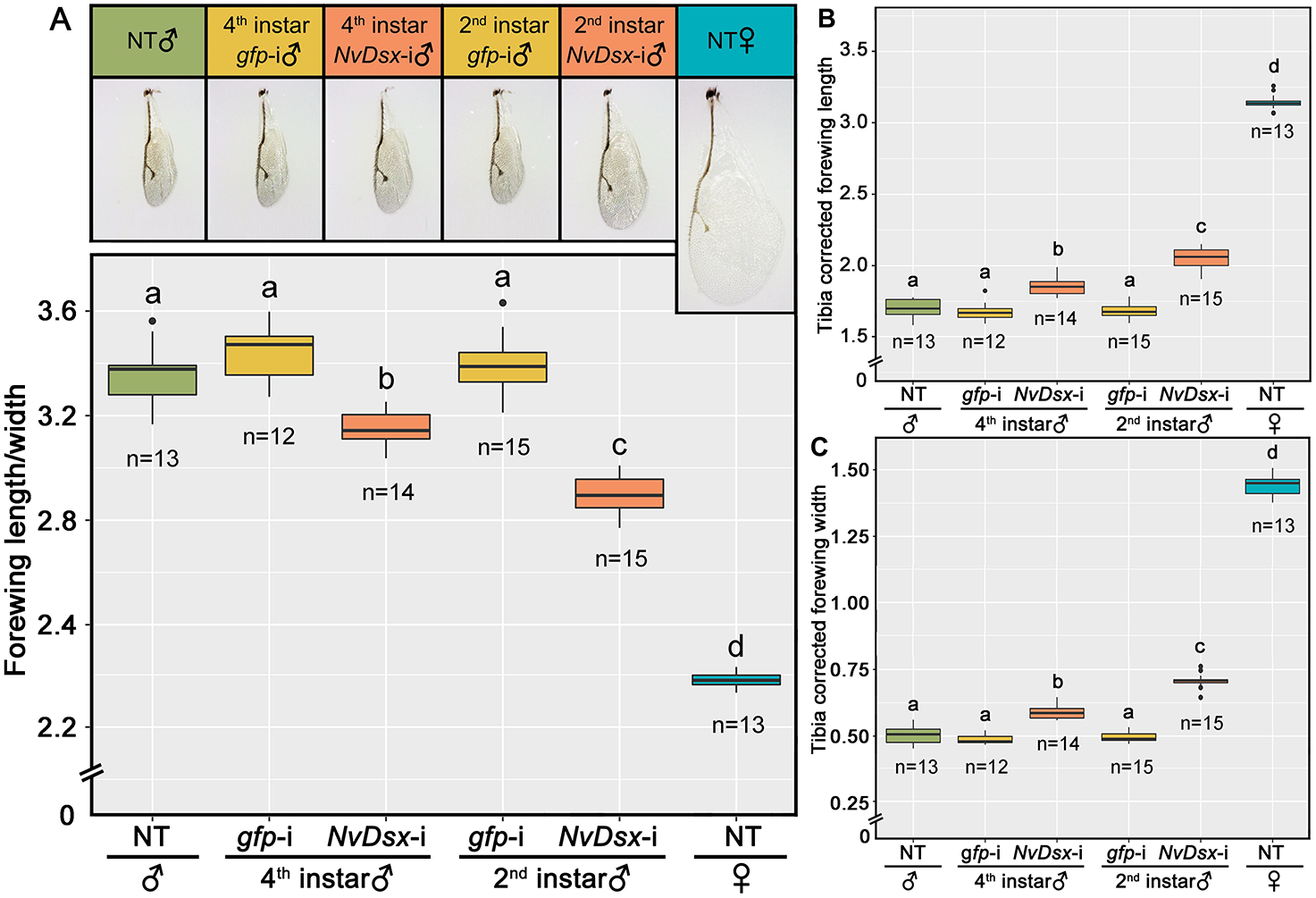
Comparison of forewing size of adult after *NvDsx* or *gfp* dsRNA treatment in the 2^nd^ or 4^th^ larval instar males and non-treated control. Y-axis represents: (A) Forewing length/width ratio corrected with tibia length, (B) Length corrected with tibia length, (C) Width corrected with tibia length. X-axis shows six different groups: males treated with dsRNA of *NvDsx* (*NvDsx*-i) and *gfp* (*gfp*-i) in the 2^nd^ and 4^th^ larval instar, non-treated (NT) males and NT females. Boxplots were generated in R and display minimum, first quartile (Q1), median, third quartile (Q3), and maximum. Dots outside the boxplots represent outliers. Replicates are provided in the figures of each treatment. One-way ANOVA and post-hoc Tukey’s HSD were used for data analysis. Letters in the figures denote statistical significance (P<0.05) among treatments.

Previous research showed that a non-coding region of *Dsx* in *N. giraulti*, termed wing-size1 locus (*ws1*), is correlated with the development of long wings in males, whereas the *ws1* variant in *N. vitripennis* is correlated with the severely reduced forewing size in males (34). Although higher *NvDsxM1* expression levels were detected in individuals with the *N. vitripennis ws1* variant (34), the mechanism of *NvDsx* regulation is unclear and the *NvDsxM2* and −*3* splice variants were unknown at the time. Our research confirmed that the regulation of the male forewing size depends on high *NvDsxM* levels and illustrates the role of *NvDsxM* in repressing wing growth. Changes in spatial and temporal expression of *unpaired-like* (*upd-like*) has been shown to affect wing width in *Nasonia* males which is caused by multiple regulatory changes (35). *NvDsxM* regulation is likely responsible for differentially expressed *upd-like*. In contrast to the observed complete female-specific pigmentation after *NvDsxM* depletion in 4^th^ instar males, the depletion of *NvDsxM* from 2^nd^ instar onwards is inadequate to restore female forewing size. All these results suggest that forewing size and shape development might already take place during embryonal development and our chosen time window for RNAi is not early enough to induce a full switch. We cannot rule out antagonistic effects of NvDsxM and NvDsxF on wing size development in *Nasonia*, since in females *Aedes aegypti, Dsx* knockdown leads to smaller wing development (36). However, we observed no wing size change after *NvDsx* knockdown in females (Fig. S5).

### Growth of male genitalia is controlled by *NvDsx*

Next, we assessed the role of *NvDsx* in male genitalia development and growth after embryogenesis is completed by depleting *NvDsx* at either 2^nd^ or 4^th^ instar larval stage. This is in contrast to many studies using genome editing in which *Dsx* is knocked out from early embryogenesis onwards (10,12). *NvDsx*-i in both developmental stages reduced the aedeagus length significantly when compared to *gfp*-i in both developmental stages and NT males (Tukey’s HSD, P<0.001) (Fig. 5A). *NvDsx*-i males from both developmental stages had the same extent of aedeagus length reduction (Tukey’s HSD, P=0.998). Aside from the shortened aedeagus, the phallomere structure (genital phallic lobes) was also severely reduced in size in both 2^nd^ and 4^th^ instar *NvDsx* depleted males (Fig. 5B-c and e) compared to *gfp*-i and NT males (Fig. 5B-a, b and d), whereas the structure of the paramere including the outer parameres, cuspis and digitus, was still intact (Fig. 5B). Compared to *gfp*-i males (Movie S1), the severe length reduction of the aedeagus rendered *NvDsx*-i males unable to copulate with females (Movie S2), showing that in this case size matters.

**Figure 5.**
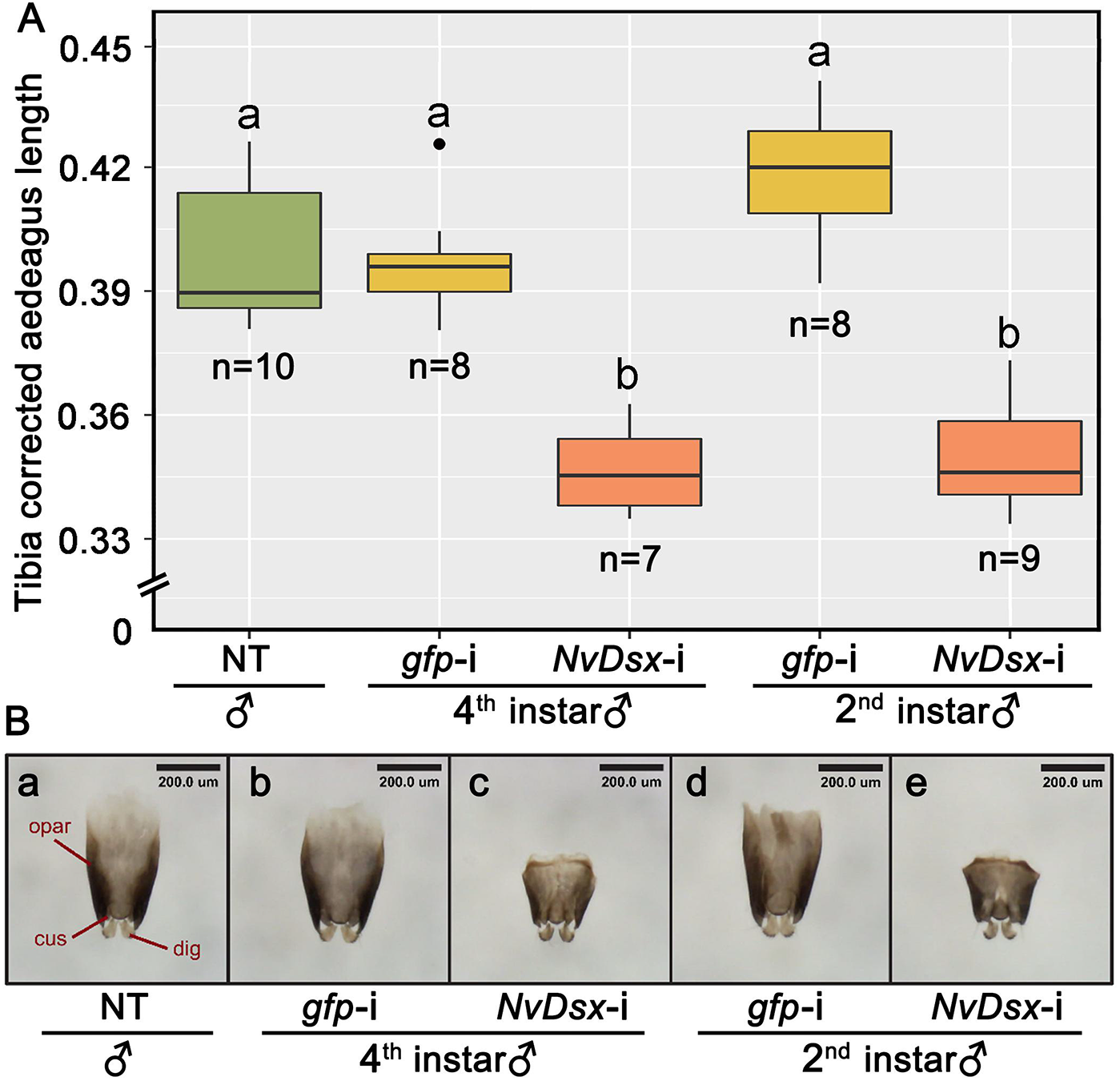
Comparison of external reproduction organs in males of different treatments. (A) Comparison of genitalia length of adult males treated with dsRNA of *NvDsx* (*NvDsx*-i) or *gfp* (*gfp*-i) in the 2^nd^ or 4^th^ larval instar males and non-treated (NT) control males. Y-axis represents genitalia length corrected with tibia length. Boxplots were generated in R and display minimum, first quartile (Q1), median, third quartile (Q3), and maximum. Dots outside the boxplot represent outliers. One-way ANOVA and post hoc Tukey’s HSD were used for data analysis. Letters in the figures denote statistical significance (P<0.05) among treatments. (B) Comparison of paramere length (without aedeagus) of adult males after *NvDsx* or *gfp* dsRNA treatment in the 2^nd^ or 4^th^ larval instar and NT control. (a) paramere structure (without aedeagus) in NT males, (b and d) 4^th^ and 2^nd^ larval instar *gfp*-i males, (c and e) 4^th^ and 2^nd^ larval instar *NvDsx*-i males. Each paramere component is described in *E. Snodgrass (1957)* (37) and labeled with the following abbreviations: outer paramere (opar), cuspis (cus) and digitus (dig).

Dsx is shown to be involved in the differentiation of genitalia structures in male and female insects in many cases by interactions with *HOX* genes during early embryogenesis (38). However, whether *Dsx* is continuously required for genitalia development has not received much attention. Previous studies using *Dsx* RNAi experiments in other species conducted at late larval stages resulted in either deformed external genitalia or chimeras that contain the structures of both sexes (24,39). Here, even though *NvDsx* RNAi was done at a distinctly earlier stage (2^nd^ larval instar) compared to other studied insect species, we did not observe any abnormalities in genital organs other than size. This shows that contrary to other studied insects, in *N. vitripennis* the differentiation of external genital organs is completed before the 2^nd^ larval instar and only growth is required during later development. It also suggests that besides the sex-specific tissue differentiation, *NvDsx* is required for the direct sex-specific trait growth as it has been shown in stag beetles *Cyclommatus metallifer* where it was demonstrated that *Dsx* can manipulate the sensitivity of JH in a sex-specific way to regulate mandible growth in males and females (14).

### Evolution of Dsx sex-specific OD2 domains and C-termini

Aligning the OD2 domains of NvDsxM1, NvDsxM2 and NvDsxM3 (Fig. S6A) or NvDsxF (Fig. S6B) with other DsxM or DsxF OD2 domains, respectively, showed that across insect orders the male-specific OD2 domain is far less conserved than the female-specific OD2 domain. Within insect orders, also a strong reduction of the sequence identity in male-specific OD2 domain alignment was seen. This observed rapid evolution of male-specific *Dsx* regions is supported by other research and is indicative of its fast evolution likely due to increased diversity of secondary sexual traits in male insects (40). In females, the sex-specific region of OD2 is relatively conserved across different insect orders. However, within the Hymenoptera, the female-specific OD2 domain showed a lower similarity. Only one out of eight conserved amino acids is being shared with all other tested species in the hymenopteran sequences, when compared to other insect orders that have half of the amino acids conserved (Fig. S6B). In *D. melanogaster* the C-terminus including the OD2 domain of DsxF binds the co-factor Intersex (41). The differences in female-specific OD2 amino acid composition in the Hymenoptera could indicate that either no co-factors or different co-factors play a role in regulating sexual development in this order compared to other insect orders. In addition, the female specific OD2 domain in the Hymenoptera was found to evolve much faster than the male-specific regions (40). This is contrary to Diptera, Lepidoptera and Coleoptera and has been suggested to correlate with the evolution of female-dominated eusociality and female-specific caste differentiation (40). However, *Nasonia* is solitary as are many other hymenopteran species. In the Hymenoptera, females arise from fertilized eggs while male development initiates by default from haploid, unfertilized eggs. We suggest that the rapid evolution of the female-specific OD2 domain in the Hymenoptera could be linked to this difference. All in all, the identification of the two additional male NvDsx isoforms opens the door to better comparative phylogenetic studies, and to better understand the evolution of *Dsx* in insects.

### Outlook

In this research we identified two sexually dimorphic traits and one male-specific trait that are actively regulated during specific developmental windows by Dsx in *N. vitripennis.* Our results showed that male-specific lack of antennal and femur pigmentation is strictly regulated during pupal development as silencing *NvDsx* from the larval stage onwards resulted in build-up of pigmentation in the males, while silencing *NvDsx* from the pupal stage onwards did not have any effect. In addition, our results suggest that sex-specific wing size and shape is regulated by *NvDsx* during embryogenesis and early larval development. As silencing *NvDsx* from the 2^nd^ larval instar onwards significantly increases wing size in males, while silencing from the pupa stage onwards does not change wing size and shape. However, *NvDsx* RNAi in the 2^nd^ larval instar did not result in complete female-like wing size, indicating that the initial wing development occurs during embryogenesis and possibly the 1^st^ larval instar. We additionally show that *NvDsx* regulates only male external reproductive organ growth and not morphology from 2^nd^ larval instar onwards, suggesting that the differentiation of external reproductive organ tissue takes place before the 2^nd^ instar which has not been reported before. Visualizing *Dsx*-expressing cells in *D. melanogaster* has demonstrated that *Dsx* gene expression is under elaborate temporal and spatial control to regulate tissue-specific sexual differentation (19). Here, we expand this knowledge using a non-drosophilid species by showing that different morphological characteristics require their own specific timing and action of Dsx protein and we speculate that this timing and action is also species-specific. Our findings solidify the view that in insects, sexual development is not only a set and forget mechanism that only operates during embryogenesis, but one that requires continuous input from Dsx on a species-specific basis.

## Material and methods

### Insects rearing

The *Nasonia vitripennis* lab strain AsymCx was reared on *Calliphora sp.* hosts *at* 16h/8h light/dark and 25°C. AsymCx strain is in the lab for over 10 years and cured from *Wolbachia* infection.

### 5’ and 3’ RACE-PCR to identify additional *NvDsx* splice variants

For identification of the 3’ ends of *NvDsx* mRNA, total RNA of 3 adult male- and female AsymCx wasps were extracted individually using TriZol (Invitrogen, Carlsbad, CA, USA) and cleared from genomic DNA by using the DNA-*free* DNA removal kit according to manufacturer’s instructions (Invitrogen, Carlsbad, CA, USA). Total RNA was annealed with the 3’ adapter from the FirstChoice RLM-RACE kit (Ambion, Austin, TX, USA) during the reverse-transcription using RevertAidTM H Minus First Strand cDNA Synthesis Kit (Fermentas, Hanover, MD, USA) and stored at −80°C. The cDNA was subjected to series of (semi-) nested PCRs using primers complementary to the adapter and gene-specific primers targeting NvDsxU, and NvDsx_X7, NvDsx_X8 and NvDsx_X9 at 94°C for 3 minutes, 35 cycles of 94°C for 30 seconds, 55-62°C for 30 seconds and 72°C for 2 minutes, with a final extension of 10 minutes at 72°C. Primers were designed based on the initial *NvDsx* gene structure (27) and sequences are listed in the Table S1.

For identification on the 5’ end of *NvDsx*, total RNA of 3 female AsymCx wasps were extracted using TriZol (Invitrogen, Carlsbad, CA, USA). After decapping, a 45-base RNA adapter was ligated to the RNA population using the FirstChoice RLM-RACE kit (Ambion, Austin, TX, USA) and reverse transcribed using the RevertAidTM H Minus First Strand cDNA Synthesis Kit (Fermentas, Hanover, MD, USA). The cDNA was subjected to series of (semi-) nested PCRs using primers complementary to the adapter and gene-specific primers targeting *NvDsxU* at 94°C for 3 minutes, 35 cycles of 94°C for 30 seconds, 55-60°C for 30 seconds and 72°C for 2 minutes, with a final extension of 10 minutes at 72°C. Primers were designed based on the initial *NvDsx* gene structure (27) and sequences are listed in the Table S1.

5’ and 3’ RACE-PCR fragments were run and visualized on ethidium bromide-containing 1.5% agarose gel with 1×TAE buffer. All RACE-PCR products were purified using GeneJET Gel Purification Kit (Fermentas, Hanover, MD, USA) and ligated into pGEM-T vector (Promega, Madison, WI, USA) and transformed into competent JM-109 *Escherichia coli* cells (Promega, Madison, WI, USA). Colony-PCRs were conducted by using primers recommended by the kit at 94°C for 3 minutes, 40 cycles of 94°C for 30 seconds, 55°C for 30 seconds and 72°C for 2 minutes, with a final extension of 7 minutes at 72°C. All isoforms were sequenced on an ABI 3730XL capillary sequencer (Applied Biosystems) and reads were visualized and aligned in MEGA4 (42). Exon-intron structure of the genes was constructed by aligning the mRNA sequences to *N. vitripennis* genome and visualizing in Geneious Prime 2019.1.3 and CLC Workbench 6.

### RT-PCR to confirm usage of alternative 1’ exon

Total RNA was extracted from four female samples and three male samples containing three adults each using NucleoSpin XS (Brand) with final elution done in 15 μl nuclease-free water. Eleven μl of eluted total RNA was converted to cDNA using RevertAidTM H Minus First Strand cDNA Synthesis Kit (Fermentas, Hanover, MD, USA) with random hexamers according to manufacturer’s protocol and stored at −80°C. One female sample was not converted to cDNA but used as gDNA contamination control in RT-PCR. The primer sets for the RT-PCR reactions are: NvEF-1a and Nv_DsxU_F6 or Nv_DsxU_F7 with Nv_Dsx_qPCR_R (Table S1). GoTaq® G2 Flexi DNA Polymerase (Promega) was used to prepare the mastermix following manufacturer’s instruction and amplified with a standard PCR profile: 3 minutes at 95°C, 35 amplification cycles of 30 seconds at 95°C, 30 seconds at 55°C, 30 second at 72°C and a final extension of 7 minutes at 72 °C in a thermal cycler (T100TM Thermal Cycler, Bio-Rad). Five μl of PCR product was visualized on a non-denaturing 1% agarose gel with a 100bp ladder. PCR fragments were Sanger sequenced at Eurofins Scientific to confirm the correct amplification.

### Alignment and phylogenetic analysis of Dsx OD2 domain across insect orders

Dsx isoforms from 22 species across insect taxa were downloaded from NCBI protein database with accession number attached in the end of each sequence name (Fig. S6A and S6B). OD2 common domain and sex-specific region of OD2 domain selection is based on An *et al*. 1996 (2). Both the male and female Dsx OD2 domain sequences were separately aligned using Guidance2 Online Server with Guidance2 method, settings are MAFFT algorithm, max iteration of 10, 100 alternative guide-trees. Geneious 2019.1 (created by Biomatters. Available from https://www.geneious.com) was used to shade the amino acids with Blosum62 score matrix applied to display the similarity between each sequence (threshold at 3).

### RNAi of *NvTraF* and *NvDsx* gene

For the *Nvtra* RNAi knockdowns, MEGAscript RNAi Kit (Thermo Fisher) was used to generate dsRNA to target exon 5-8 of *Nvtra* mRNA in 4th larval instar females which resulted in a male splice form of *NvDsx* as shown in Verhulst *et al.* (28). To conduct functional analysis of *NvDsx*, same RNAi Kit was used to generate dsRNA based on either the common region of *NvDsx* (exon 2-5) that presents in both male and female splice-variants or the specific region of *NvDsx* (exon 7-9) that only presents in two longer male splice-variants. *Gfp* dsRNA was generated from the vector pOPINEneo-3C-GFP which was a gift from Ray Owens (Addgene plasmid # 53534; http://n2t.net/addgene: 53534; RRID: Addgene_53534). This *gfp* dsRNA was used as control in all experiments as it has no target in *Nasonia*. Primers designed using Geneious 10.0.9 to construct the dsRNA T7 template are listed in the Table S1. For male-specific *NvDsx* RNAi, offspring of AsymCX strain was collected at 2^nd^ and 4^th^ larval instars and white pupa stage which are described in Table S2. All dsRNAs were diluted with RNase-free elution buffer (MEGAscript RNAi Kit (Thermo Fisher)) to 4000 ng/μl (NanoDrop™ 2000 Spectrophotometer, ThermoFisher) before proceeding to microinjection.

Microinjection of wasps was carried out with IM300 Microinjecctor (Narishige) and FemtoJet® 4i (eppendorf) and followed the protocols described by Werren *et al*. (43) and Lynch & Desplan (44) with minor changes. Red colour food dye was pre-mixed with the dsRNAs in a ratio of 1:9. Experiments were performed using AsymCx strain at different developmental stages for different purposes (Table S3).

Second and 4^th^ instar larvae were collected from parasitoid hosts and placed on a 1X PBS agar plate before injection. Early pupal stage samples were fixed on the glass slides with double-sided tape. After injection, 2^nd^ instar larvae were transferred back to host (6-8 per host) and sealed with the host shell. Afterward, sealed hosts were kept in 8 tube-PCR strips with open lid. Subsequently, all injected samples either in 8 tube-PCR strip or on slides were placed on 1X PBS plates to prevent dehydration and incubated at standard rearing conditions until adult emergence.

### *NvDsx* splicing and expression throughout development

Male and female wasps were separated at black pupal stage by the sex-specific traits such as forewing size and presence/absence of the ovipositor. To generate female offspring, one selected male and one selected female pupa were collected in a single glass vial clogged with cotton plug until eclosion and kept for one more day to make sure they had mated. After fertilization, two hosts per day for the first two days were provided to mated females to initiate oviposition. Afterwards, one-hour host presentation was repeated twice in a row per day. Since mated *N. vitripennis* tend to lay extremely female-biased eggs, offspring from mated females was considered female (45). Male offspring was collected by using virgin females with the same setup. To facilitate early stage (eggs) sample collection, egg-laying chambers (1.5 ml Eppendorf tubes with cut-off tip presenting the head of host pupa plugged in 1000 μL filter pipet tips) were made to allow the females only access to the head portion of the fly hosts.

To confirm that male-specific splice variants are not under the regulation of NvTraF but splice in default mode independently, female samples were collected from 3h, 1d, 2d, 3d, 4d, 5d and 6d after Nv*traF* knockdown. To study the expression pattern of *NvDsx* sex-specific splice variants throughout whole developmental stages, samples were collected at specific time points listed in Table S2. More than 50 embryo eggs were collected at 36h and five to 10 larvae, pupae or adults were collected at 2.5d and onwards. Collected samples were subsequently frozen in liquid nitrogen and stored in −80°C. Sex-specific primers of *NvDsx* were designed using Geneious 10.0.9. and are listed in the Table S1. Both *NvRP49* and *NvEF-1a* were assessed for their expression stability in this semi-quantitative RT-PCR setup, and *NvEF-1a* was finally chosen as the reference gene.

Frozen samples were homogenized by pestles which were designed for 1.5 ml microcentrifuge tubes (Biosphere SafeSeal Tube 1.5ml). Total RNA was extracted using ZR Tissue & Insect RNA MicroPrep™(Zymo) following manufacturer’s instructions. On column DNase treatment step was added to all samples. Subsequently, total RNA of each samples was eluted in 16 μl of DNase/RNase free water. After verifying the purity and concentration by NanoDrop™ 2000 Spectrophotometer (ThermoFisher), one μg of each template was synthesized into cDNA with a standard reaction mix (SensiFAST™ cDNA Synthesis Kit, Bioline) in a thermal cycler (Bio-Rad T100TM Thermal Cycler, Bio-Rad) with 5 minutes priming at 25°C, 30 minutes reverse transcription at 46°C and 5 minutes reverse transcriptase inactivation at 85°C.

For all RT-PCR reactions, GoTaq® G2 Flexi DNA Polymerase (Promega) was used to prepare the mastermix following manufacture’s instruction and amplified with a standard PCR profile: 3 minutes at 95°C, 34 (male-specific splicing verification) / 29 (expression pattern of *NvDsx*) amplification cycles of 30 seconds at 95°C, 30 seconds at 55°C, 50 seconds at 72°C and a final extension of 5 minutes at 72 °C in a thermal cycler (T100TM Thermal Cycler, Bio-Rad). In order to standardize the expression pattern of *NvDsx*, the number of PCR cycles needed was determined by comparison of the brightness of the bands in gels from different PCR reactions with cycles ranging from 25 to 34 to achieve non-oversaturated brightness. In the end 29 cycles were used for both reference genes and target genes.

### Silencing efficiency of *NvDsx*

qPCR was conducted to verify the silencing efficiency of *Dsx* knockdown. After *dsRNA* injection, samples were collected at either adult stages (injected at the 4^th^ larval instar and the white pupa stage) or the white pupa stage (injected at the 2^nd^ larval instar). Four to five pupae or adults were pooled to produce one biological replicate and six biological replicates per treatment were prepared. All testing samples were used in qPCR according to SensiFASTTM SYBR® No-ROX Kit manual (Bioline). *NvDsx* qPCR primers were designed on the common region (exon 2-5) of male and female splice variants (Table S1). *NvEF-1a* was used as reference genes. qPCR was carried out using the CFX96TM Real-Time System (Bio-Rad) with Bio-Rad CFX Manager 3.1 Software (Bio-Rad). The standard qPCR profile consists of 95°C for 3 minutes, 45 amplification cycles of 15 seconds at 95,15 seconds of 55°C, 30 seconds of 72°C and a final standard dissociation curve step to check for non-specific amplification.

### Measurements of male forewing size, external reproduction organs

Forewing size was measured by wing length and width based on the standard methods described by Loehlin *et al.* (46). Tibia length of the hind leg was used to standardize the body size differences among individuals (47). Forewings and external reproduction organ of adult males were dissected on clean slides with a drop of 1 X PBS buffer using dissection tweezers and subsequently mounted in a drop of ethanol on slides which was allowed to evaporate. External reproduction apparatuses were identified based on the description in Liu *et al*. (48) and E. Snodgrass (37). Photos were taken by Dino-Lite Edge 5MP digital microscope and measured by using DinoCapture 2.0 software.

### Data analysis

qPCR data was first imported to LinRegPCR software (LinRegPCR, 2017.1.0.0, HFRC, Amsterdam, The Netherlands) (49). After baseline correction, the initial number of templates (N0) were calculated based on the average PCR efficiency of each amplicon. Relative expression levels of *NvDsx* in each sample was obtained by dividing the N0 value of *NvDsx* by N0 value of *NvEF-1a*. Statistical analysis of qPCR and measurement data was performed in R using One-way ANOVA and Tukey’s Honest Significant Difference (HSD) for post hoc test.

## Acknowledgment

We thank Hans Smid for assisting with phenotype recording; Rutger Diepeveen for his contribution to the *NvDsx* RNAi work; Joan Diaz Calafat, Age Muller, Simon Pleiter and Romy Gielings for their contribution to the *NvtraF* RNAi work; Min Xu for supporting the sample preparation, collection and measuring; and Weizhao Sun for recording the mating behavior of RNAi treated males. We additionally thank all the tested *N. vitripennis* for their great sacrifice to science. This work is part of the Dutch Research Council (NWO) research programme Innovational Research Incentives Scheme Veni with project number 863.13.014 granted to ECV.

---

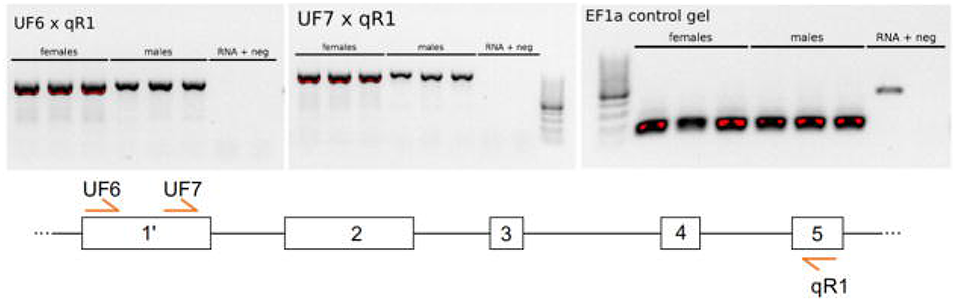

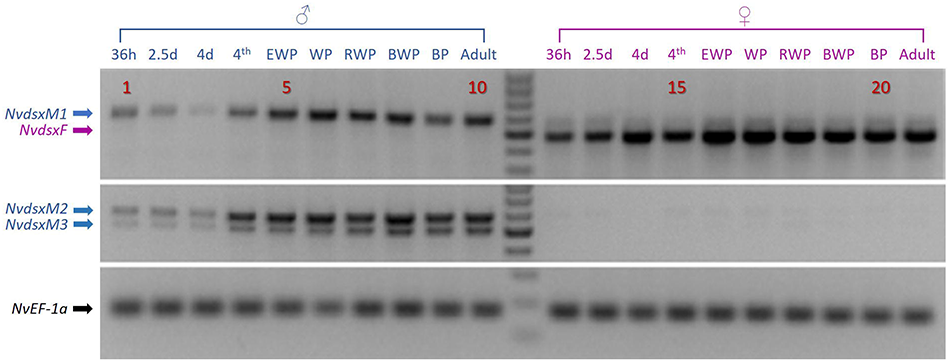

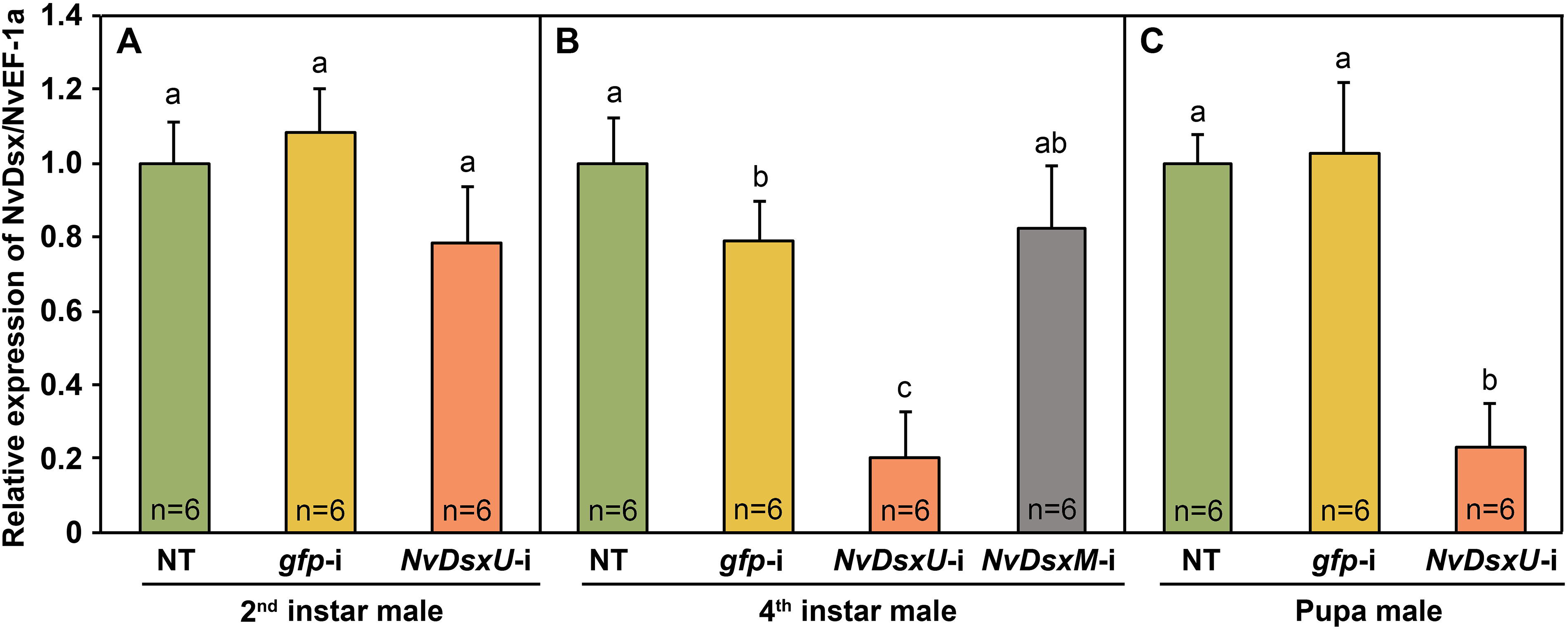

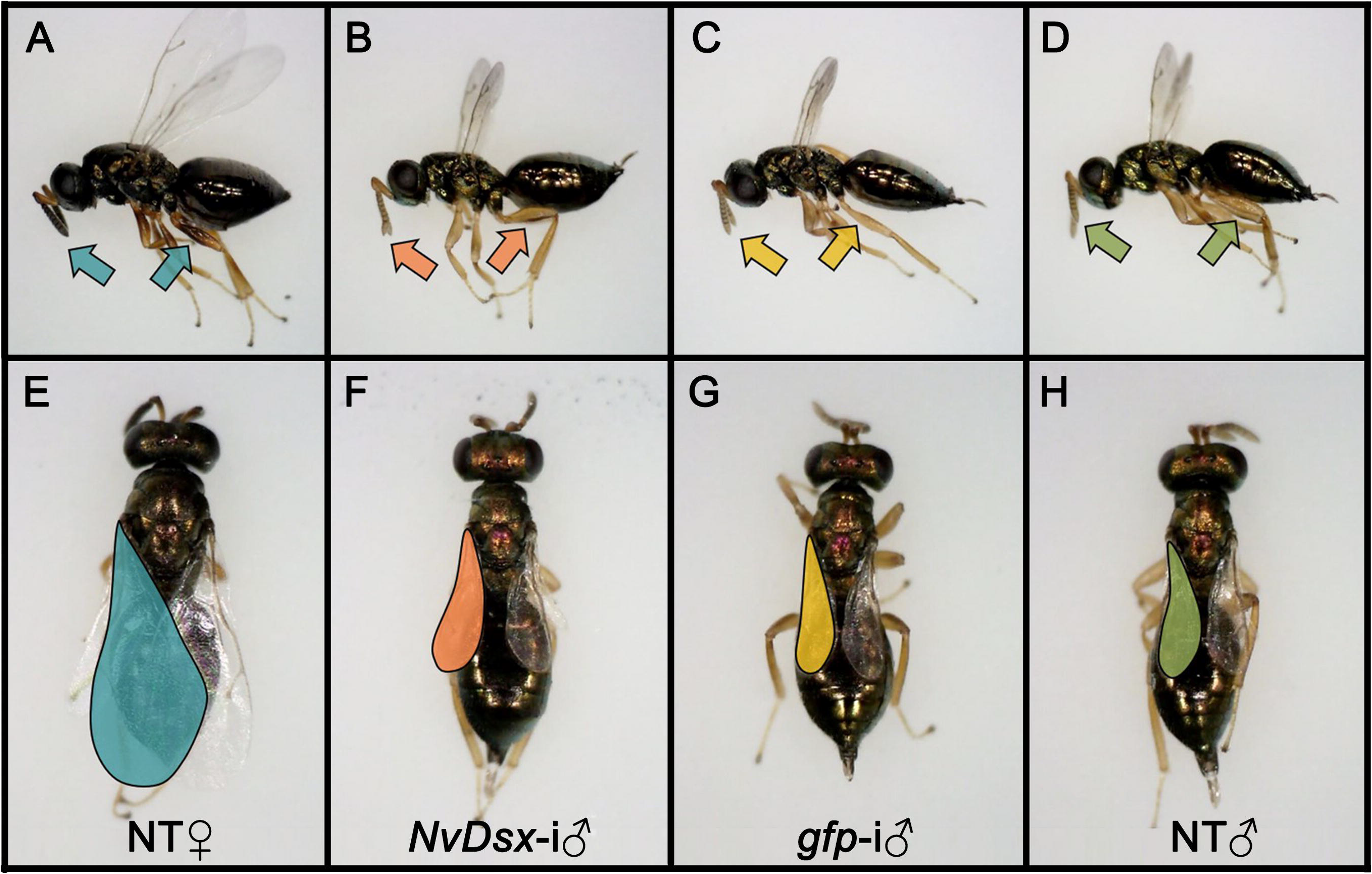

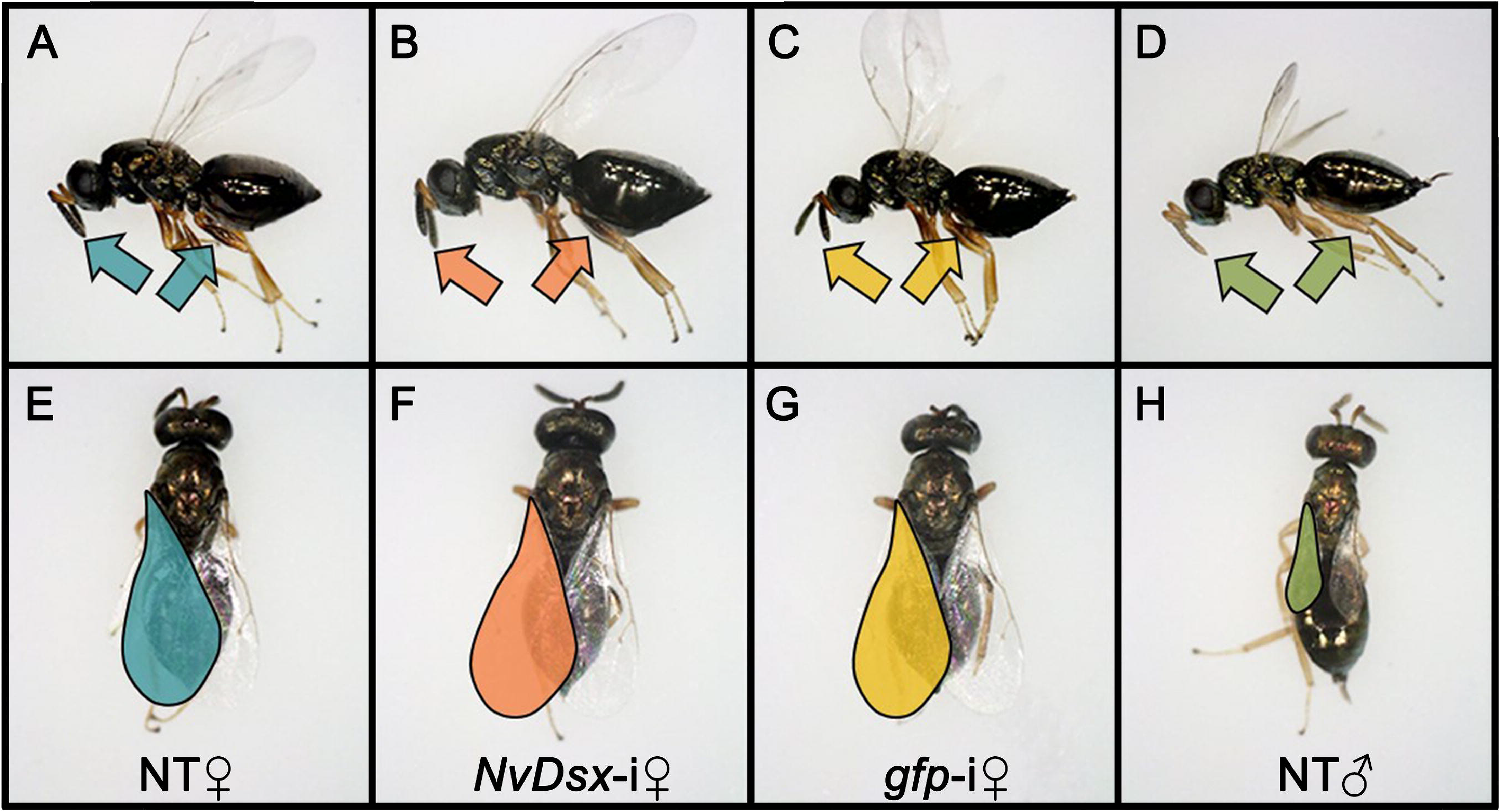

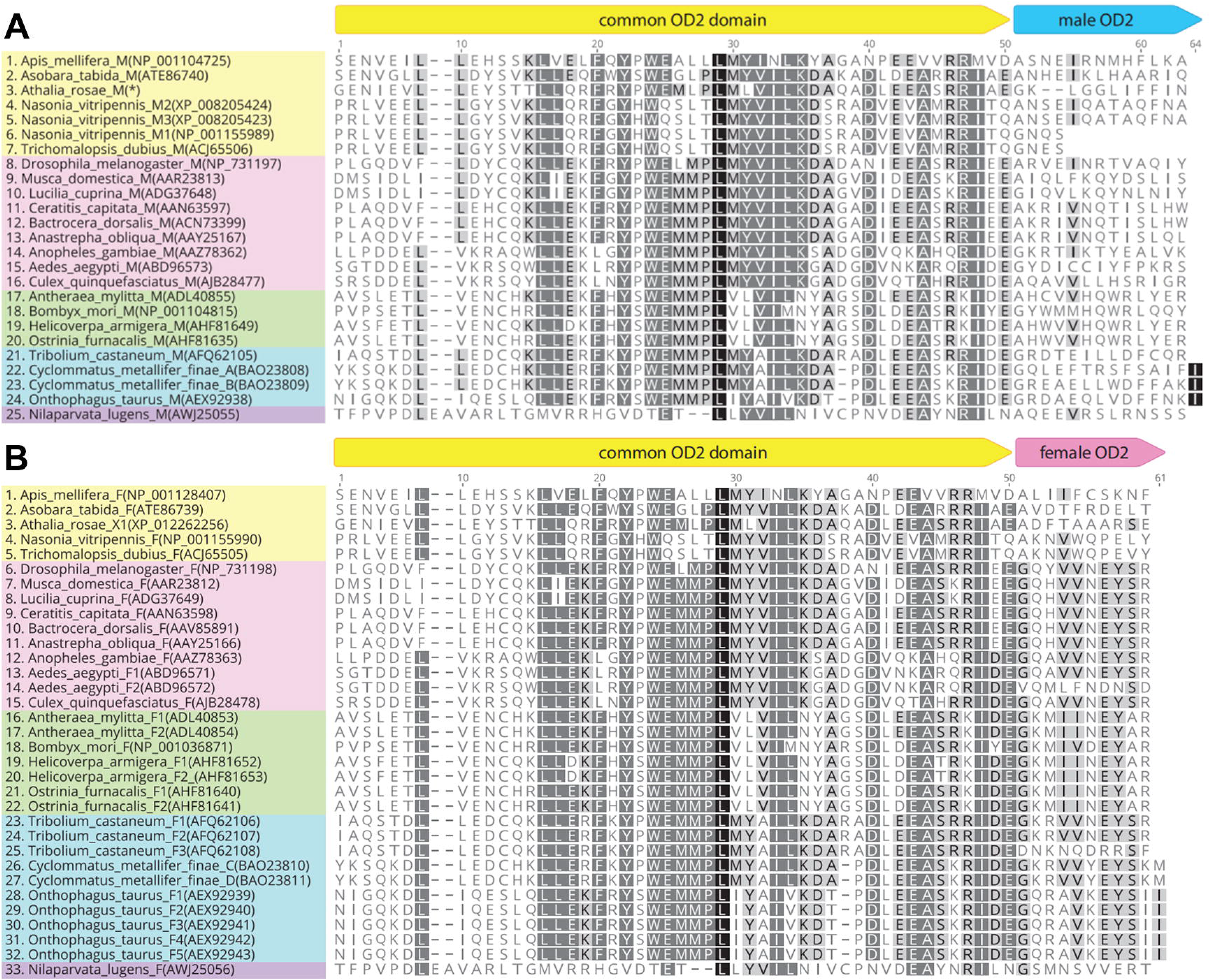

## Notes

https://www.dx.doi.org/10.6084/m9.figshare.12152322

### Competing Interest Statement

The authors have declared no competing interest.

### Summary of Updates

V2: Introduction clarified; textual errors removed; author affiliations updated; full size images added. V3: Legend mistake for Figure S4 fixed.

