## Supplementary information for "Sexually dimorphic traits and male-specific differentiation are actively regulated by Doublesex during specific developmental windows in *Nasonia vitripennis*"

**This PDF file includes:**

Supplementary text

Figures S1 to S6

Tables S1 to S3

Legends for Movies S1 to S2

SI References

**Other supplementary materials for this manuscript include the following:**

Movies S1 to S2

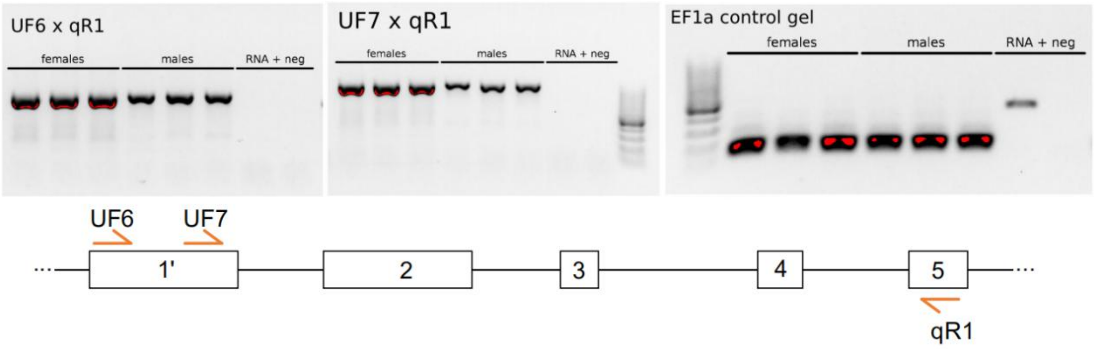


**Fig. S1. Schematic overview of putative *NvDsx* exons with the correlated RT-PCR results.** RT-PCR shows the joining of exon 1’ with exon 2 of the *NvDsx* transcript. A higher usage of exon 1’ in females compared to males is observed. *NvEF-1a* was used as expression control. RNA indicates sample that has not been converted to cDNA and includes any remaining gDNA in the sample. Neg. is negative template RT-PCR control. Primer qR1 is Nv_Dsx_qPCR_R (Table S1). All samples were loaded on 1% agar gel with 100bp ladder (Thermo).

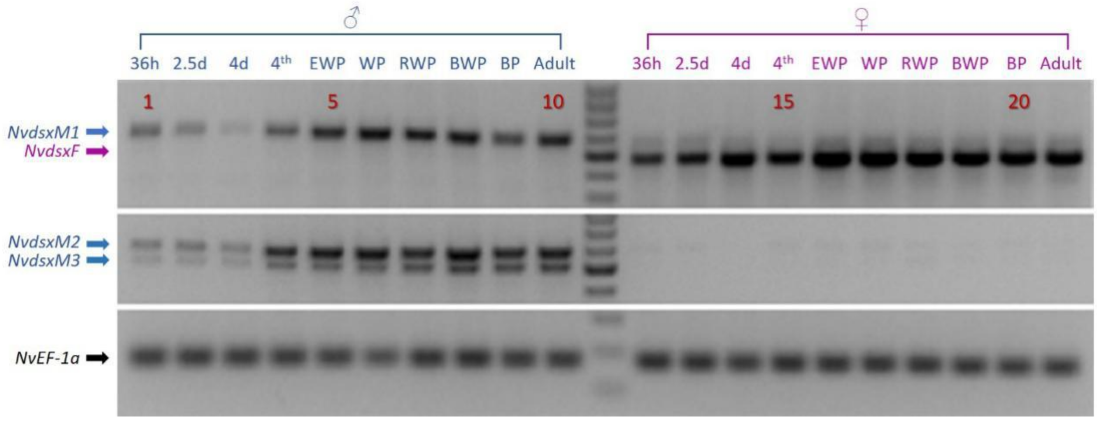


**Fig. S2. Confirmation of sex-specific *NvDsx* transcript expression throughout different developmental stages in *N. vitripennis* males and females.** The overall expression profile of each splice variant throughout ten developmental stages ranging from embryo to adult was determined (Table S2). Arrows indicate male (in blue) and female (in purple) splice forms. Lanes 1-10: *N. vitripennis* male samples from 36h to adult stage (2.5d: 2.5 day, 4d: 4 day, 4th: 4$^{th}$ larval instar, EWP: early white pupa stage, WP: white pupa stage, RWP: red eye white pupa stage, BWP: black and white pupa stage, BP: black pupa stage, Adult: Adult stage). Lanes 11: 100 bp ladder. Lanes 12-21: *N. vitripennis* female samples from 36h to adult stage. All samples were loaded on 1.5% agar gel with 100bp ladder (Thermo) indication. *NvEF-1a* was used as an expression control.

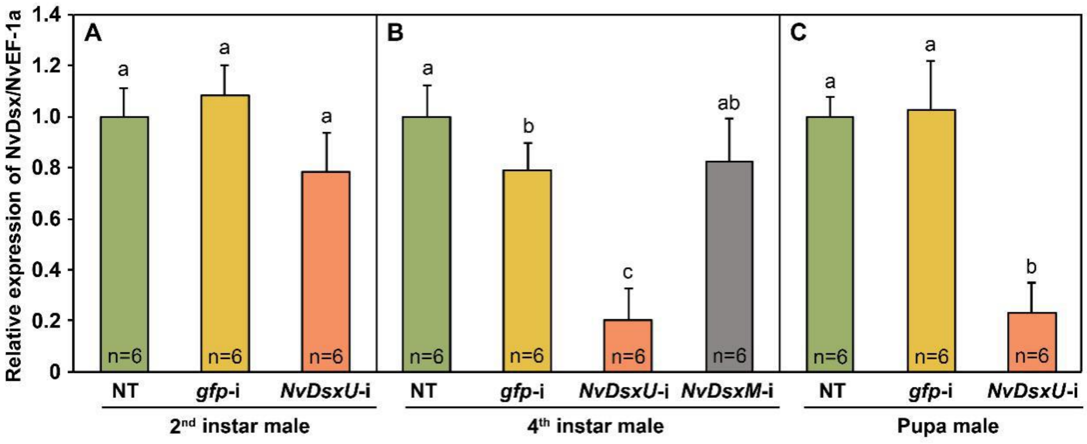


**Fig. S3. Relative expression levels of *NvDsx* after RNAi in different developmental stages.** Relative expression of *NvDsx* / *NvEF-1a* among non-treated and second (A) and fourth instar treated males (B) and white pupa stage treated males (C). To verify the function of *NvDsxM* in general and *NvDsxM2* and *NvDsxM3* specifically, dsRNAs NvDsxU and NvDsxM were designed against the common region of *NvDsx* (exon 2-5) and the common region of *NvDsxM2* and *NvDsxM3* (exon 7-9) respectively. A *gfp* dsRNA was used as a mock. *NvEF-1a* was used to standardize the variance between samples. X-axis shows different treatments: *gfp*-i*,* *NvDsxU*-i and *NvDsxM*-i refers to *gfp* knockdown, *NvDsx* knockdown targeting the common region and *NvDsx* knockdown targeting the *NvDsxM* sex specific region respectively. Y-axis shows percentage of *NvDsx* / *NvEF-1a* expression of treatments to NT control. Error bars show standard error (n=6). ANOVA and post hoc Tukey's HSD were used for data analysis. Letters in the figures show statistical significance (P<0.05) among treatments.

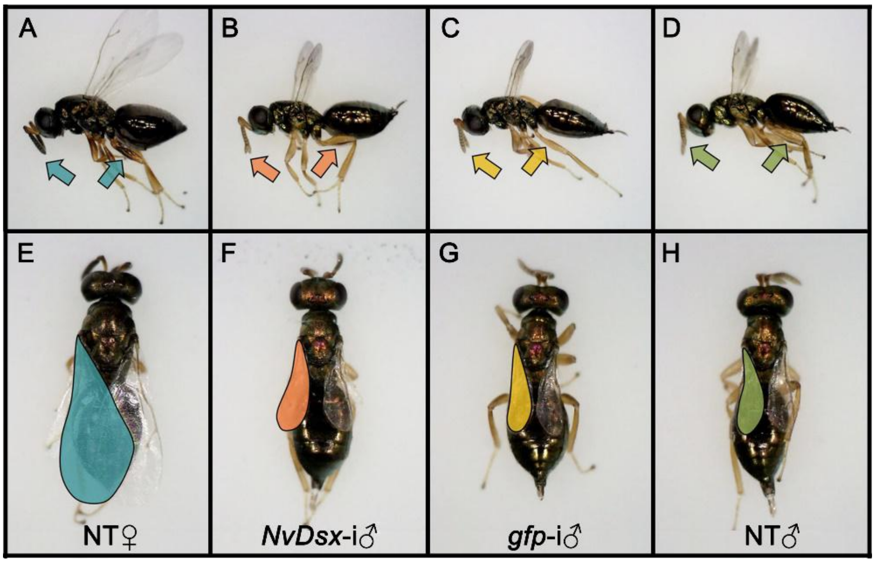


**Fig. S4. Adult phenotype comparison of early white pupal stage *NvDsx* knockdown males with *gfp* control injected males.** Comparison of early white pupal stage *NvDsx* knockdown (*NvDsx*-i) male (B and F) with mock *gfp* knockdown (*gfp*-i) male (C and G), non-treated (NT) male (D and H) and NT female (A and E). Samples were collected at adult stage and killed by freezing in liquid nitrogen before being placed on glass slides for photo shooting. Top row shows the lateral side of tested *N.vitripennis*. Two arrows in each photo indicate the position of pigmentation in the antennae and hind legs. Compared to NT males (D), none of the *NvDsx*-i males (B) and *gfp*-i males (C) have pigmentation in antennae and hind legs, while NT females have dark pigmentation in the antennae and hind legs (A). Bottom row shows the dorsal side of tested *N.vitripennis*. Colours mark forewing area of different treatments. Compared among NT males (H), *NvDsx*-i males (F) and *gfp*-i males (G), they all have relatively smaller forewing size and there is no observable difference among them, while NT females have large forewings (E).

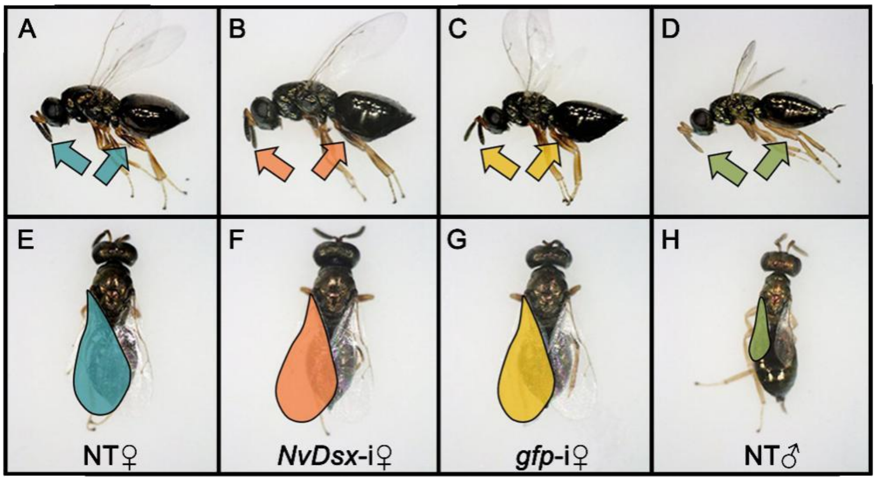


**Fig. S5. Adult phenotype comparison of 4th instar *NvDsx* knockdown females with mock *gfp* injected females.** Comparison of 4th instar *NvDsx* knockdown (*NvDsx*-i) female (B and F) with mock *gfp* knockdown (*gfp*-i) female (C and G), non-treated (NT) male (D and H) and NT female (A and E). Samples were collected at adult stage and killed by freezing in liquid nitrogen before being placed on glass slides for photo shooting. A, B, C and D show the lateral side of the tested *N.vitripennis*. Two arrows in each photo indicate the position of pigmentation in antennae and hind legs. Compared to NT females (A), none of the *NvDsx*-i or *gfp*-i females show loss of pigmentation in antennae and hind legs (B and C), while males have no pigmentation (D). E, F, G and H show the dorsal side of tested *N.vitripennis*. Colours mark forewing area of different treatments. Compared with NT females (E), *NvDsx*-i females and *gfp*-i females have relatively larger forewing size and there is no observable difference among them (F and G), while males have short and narrow forewings (H).

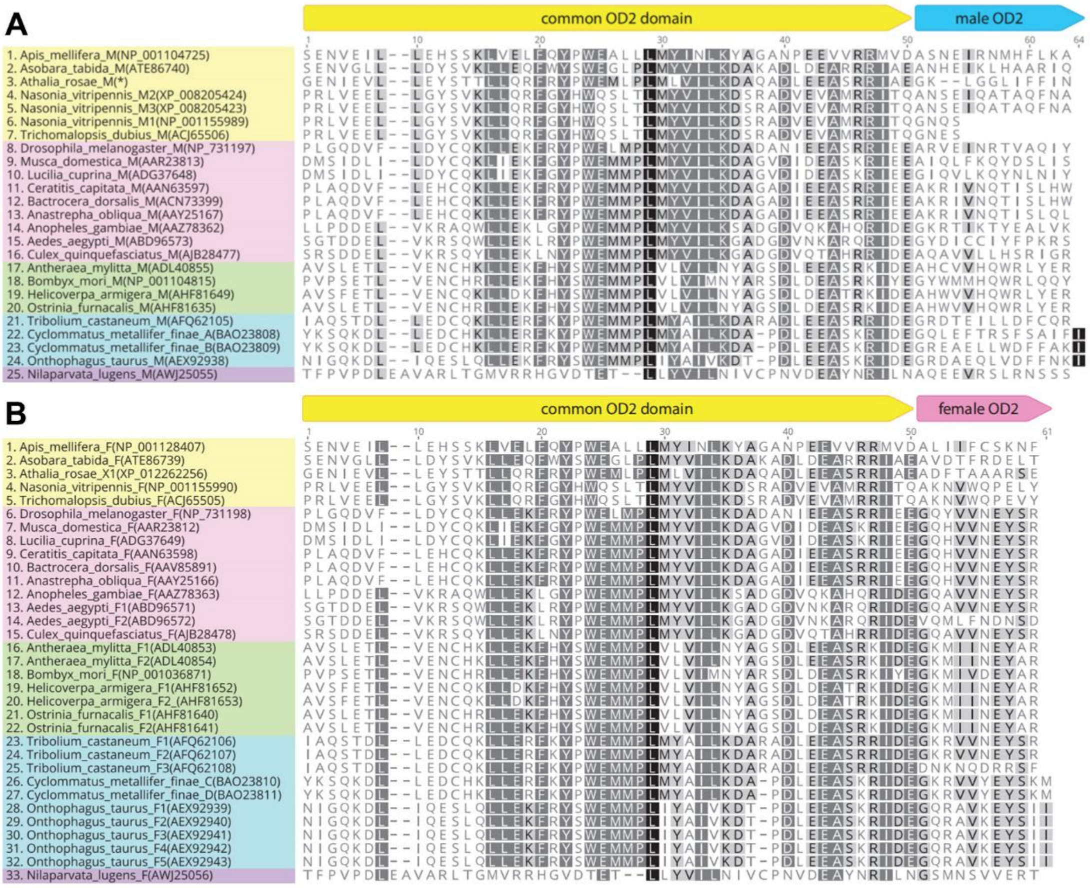


**Fig. S6. Comparison of common and sex-specific OD domain sequences across Dsx proteins in different insect orders.** Amino acid sequence alignments of common OD2 domain and male-specific (A) or female-specific (B) Dsx OD2 in species across different insect orders. Common OD2 and sex-specific regions are indicated above the figure panels. Sites which are shaded by the colour light grey, dark grey and black indicate 60-80%, 80-100% and 100% similarity among all aligned sequences. Start position of common OD2 and male OD2 is based on *D. melanogaster* (1). Species names which belong to the same insect order are in a uniform colour block, yellow represents Hymenoptera, pink represents Diptera, green represents Lepidoptera, blue represents Coleoptera and purple represents Hemiptera. Athalia_rosae_M Dsx sequence was extracted from publication Mine *et al.* (2) and is indicated with an asterisk.

**Table S1. Primers used in this research.** 'Region' indicates which *NvDsx* or *NvTra* exons are targeted from the given primer sets. * Sequences in '[]' represent the T7 adaptor sequence that provided by the MEGAscript RNAi Kit (Thermo Fisher).

a. (Verhulst *et al*., 2010) (3)

| Target | Region | Primer | Sequence |
|---|---|---|---|
| | | 3' race adapter | 5'-GCGAGCACAGAATTAATACGACTCACTATAGGT12VN-3' |
| | | 3' race outer primer | 5'-GCGAGCACAGAATTAATACGACT-3' |
| | | 3' race inner primer | 5'-CGCGGATCCGAATTAATACGACTCACTATAGG-3' |
| | | 5' race outer primer | 5'-GCTGATGGCGATGAATGAACACTG-3' |
| | | 5' race inner primer | 5'-CGCGGATCCGAACACTGCGTTTGCTGGCTTTGATG-3' |
| NvDsxM2 & NvDsxM3 | Exon 5-9 | A14_Nvit_DsxU_F1 | 5'-ACGGTTTGGCTACCATTGGC-3' |
| | | Nv_DsxM_RNAi_R | 5'-TAGTAGACGCTCTCTTTGGG-3' |
| NvDsxF & NvDsxM1 | Exon 4-6 | A14_Nvit_DsxU_F2 | 5'-TGAGCAATAACAACAGCAACAG-3' |
| | | Nv_DsxM_R1[a] | 5'-TCGGAGAAGATTGGCAGAAC-3' |
| NvDsx common region | Exon 2 | A14_Nv_DsxU_R2 | 5'-GTATGCGCTGACTTGGCTTG-3' |
| | | A14_Nv_DsxU_R3 | 5'-GACTTGGCTTGGGCTTTTTCGTC-3' |
| | | A14_Nv_DsxU_R4 | 5'-ACACTTTTGCACTGCAGACCCT-3' |
| | Exon 3-5 | Nv_Dsx_qPCR_F[a] | 5'-CAGCAACACGAGATGCTGATGG-3' |
| | | Nv_Dsx_qPCR_R[a] | 5'-TGTCATTACTGCCATTCATGCTTGG-3' |
| | Exon 1' | Nv_DsxU_F6 | 5'-CAAGCAGAGAGCAGCTCAGG-3' |
| | | Nv_DsxU_F7 | 5'-ACAACGGAGCAAGAGGGAAG-3' |
| | Exon 2-5 | Nv_DsxU_RNAi_F | 5'-*[TAATACGACTCACTATAGGG]CCAAGAGGCAGCAAATTATG-3' |
| | | Nv_DsxU_RNAi_R | 5'-*[TAATACGACTCACTATAGGG]GTTATACGCCGCATGGCTAC-3' |
| NvDsx male-specific region | Exon 7-9 | Nv_DsxM_RNAi_F | 5'-* [TAATACGACTCACTATAGGG]CGGCTACTATCCACCCTCG-3' |
| | | Nv_DsxM_RNAi_R | 5'-* [TAATACGACTCACTATAGGG]TAGTAGACGCTCTCTTTGGG-3' |
| | Exon 7 | A14_Nvit_Dsx_X7_R1 | 5'-CAAAGGCGTAGGGCAGGAG-3' |
| | Exon 8 | A14_Nvit_Dsx_X8_R1 | 5'-GACGCTTGCTTAATCCGTGG-3' |
| | Exon 9 | A14_Nvit_Dsx_X9_R1 | 5'-AATCCTCGGCCGGAAATAGC-3' |
| Nvtra common region | Exon 5-8 | Nv_Tra_RNAi_F1[a] | 5'-*[TAATACGACTCACTATAGGG]CGAGACATCAGTTAGAAGAT-3' |
| | | Nv_Tra_RNAi_R1[a] | 5'-*[TAATACGACTCACTATAGGG]GTCTTGTGGTCCTATGAAAC-3' |
| NvtraF & NvtraM | Exon 2-3 | Nv_Tra_F2[a] | 5'-GACCAAAAGAGGCACCAAAA-3' |
| | | Nv_Tra_R3[a] | 5'-GGCGCTCTTCCACTTCAAT-3' |
| NvEF-1a | | Nv_EF-1a_qPCR_F[a] | 5'-CACTTGATCTACAAATGCGG-3' |
| | | Nv_EF-1a_qPCR_R[a] | 5'-GAAGTCTCGAATTTCCACAG-3' |

**Table S2**. **Developmental stages and time points in the development of *Nasonia vitripennis*.** Overview of different developmental stages and time points used in this study with their abbreviations.

| Abbreviation | Description | Development stage | Duration of development at 25 °C, LD 16:8 |
|---|---|---|---|
| 36h | 1$^{st}$ instar larva | Embryo/larva | 36 h |
| 2.5d | 2$^{nd}$ instar larva | Larva | 2.5~3 d |
| 4d | 3$^{rd}$ instar larva | Larva | 4 d |
| 4$^{th}$ | 4$^{th}$ instar larva | Larva | ~ 6 d |
| EWP | Early white pupa | Pupa | ~ 7 d |
| WP | White pupa | Pupa | ~ 8 d |
| RWP | White pupa, red eyes | Pupa | ~ 9 d |
| BWP | Black pupa, white abdomen | Pupa | ~ 10 d |
| BP | Black pupa | Pupa | ~ 12 d |
| Adult | Adult | Imago | ~ 14 d |

**Table S3. Target genes and developmental stages of *Nasonia vitripennis* for microinjection for different purposes.**

| Purpose | Target genes | Targeted Development stages |
|---|---|---|
| ***NvDsxM*** **splicing variants verification** | ***gfp & NvtraF*** | **$4^{th}$ larval instar** |
| ***NvDsx*** **functional analysis** | ***gfp, NvDsx & NvtraF*** | **$2^{nd}$ larval instar, $4^{th}$ larval instar & early white pupa** |

**Movie S1 (separate file). A naive adult male injected with *gfp* dsRNA at the 4th instar larval stage courts with an untreated naive female adult.** Male is on top and performs a series of head nods in which pheromones are exchanged with the female. At 8 seconds the female becomes receptive and the male slides down to copulate. At 20 seconds, the male climbs back up and performs post-copulatory head nods to end the copulation.

**Movie S2 (separate file). A naive adult male injected with dsx dsRNA at the 4th instar larval stage courts an untreated naive female adult.** Male is on top and performs a series of head nods in which pheromones are exchanged with the female. At 8 seconds the female becomes receptive and the male slides down in an attempt to copulate. The male tries to copulate for 21 seconds until the female walks away at 30 seconds, and no copulation took place.

Supplementary movies available on https://www.dx.doi.org/10.6084/m9.figshare.12152322
